# Horizontal gene transfer to a defensive symbiont with a reduced genome amongst a multipartite beetle microbiome

**DOI:** 10.1101/780619

**Authors:** Samantha C. Waterworth, Laura V. Flórez, Evan R. Rees, Christian Hertweck, Martin Kaltenpoth, Jason C. Kwan

## Abstract

The loss of functions required for independent life when living within a host gives rise to reduced genomes in obligate bacterial symbionts. Although this phenomenon can be explained by existing evolutionary models, its initiation is not well understood. Here, we describe the microbiome associated with eggs of the beetle *Lagria villosa*, containing multiple bacterial symbionts related to *Burkholderia gladioli* including a reduced-genome symbiont thought to produce the defensive compound lagriamide. We find that the putative lagriamide producer is the only symbiont undergoing genome reduction, and that it has already lost most primary metabolism and DNA repair pathways. The horizontal acquisition of the lagriamide biosynthetic gene cluster likely preceded genome reduction, and unexpectedly we found that the symbiont accepted additional genes horizontally during genome reduction, even though it lacks the capacity for homologous recombination. These horizontal gene transfers suggest that absolute genetic isolation is not a requirement for genome reduction.

## INTRODUCTION

Mutualistic symbioses between animals and bacteria, widespread in nature, serve a variety of functions such as biosynthesis of nutrients not found in the host’s diet (Akman et al., 2002; Shigenobu et al., 2000), and protection from predation (Lopera et al., 2017; Miller et al., 2016a; Piel, 2002) or infection (Currie et al., 1999; Flórez et al., 2018; Kroiss et al., 2010). Such relationships exist on a continuous spectrum of dependency and exclusivity from the perspective of both the host and symbiont. Symbionts generally become obligate after a prolonged period of exclusive association with the host, and the symbionts that become obligate tend to carry out highly important functions for the host (Latorre and Manzano-Marín, 2017; Lo et al., 2016; McCutcheon and Moran, 2012). For example, mitochondria and chloroplasts, organelles that are required for energy production and carbon fixation in eukaryotic and plant cells, originated from endosymbiotic capture of alphaproteobacteria and cyanobacteria ∼1.2 Bya and ∼900 Mya, respectively (Shih and Matzke, 2013). The acquisition of these organelles allowed the diversification of eukaryotic species (López-García et al., 2017). More recently, aphids evolved to feed on plant sap depleted of several essential amino acids only through capture of an endosymbiont, *Buchnera aphidicola*, that can synthesize these nutrients, ∼160–280 Mya (Moran et al., 1993; Munson et al., 1991). In these cases, the microbial symbiont has lost the ability to live independently of the host, and the hosts are also dependent on their symbionts.

The mechanism by which symbionts become obligate is through loss of genes required for independent (but not host-associated) life, leading to an overall reduction in genome size (Latorre and Manzano-Marín, 2017; Lo et al., 2016; McCutcheon and Moran, 2012). This gene loss is the result of relaxed selection on genes for functions provided by the host, and also increased genetic drift as a result of small effective populations in strict vertical transmission (Latorre and Manzano-Marín, 2017). A general mutational bias towards deletion in bacteria (Mira et al., 2001) combined with many successive population bottlenecks that allow the fixation of slightly deleterious mutations (Latorre and Manzano-Marín, 2017) mediates general gene degradation and genome reduction. While these processes are largely thought to be nonadaptive, there is some evidence that increase in AT and reduction in genome size could be selected for to reduce metabolic burden on the host (Dietel et al., 2019, 2018). The early stages of this process are manifested by a proliferation of nonfunctional pseudogenes and a decrease in coding density in the genome (Lo et al., 2016), before the intergenic sequences are lost to eventually give tiny <1 Mbp genomes (Latorre and Manzano-Marín, 2017; McCutcheon and Moran, 2012). While there is a robust model for the evolutionary forces that drive this process once a symbiont becomes host-restricted, it is unknown how bacteria first become obligate and start on the road to genome reduction (Latorre and Manzano-Marín, 2017).

The beetle subfamily Lagriinae offers an opportunity to examine this question. Various Lagriinae have evolved special symbiont-bearing structures that serve to facilitate the vertical transmission of bacteria (Flórez et al., 2017). Beetles are typically co-infected with multiple symbiont strains related to the plant pathogen *Burkholderia gladioli*, that are secreted onto the surface of eggs as they are laid (Flórez et al., 2017). In the species *Lagria villosa*, a South American soybean pest, at least one symbiotic *B. gladioli* strain (Lv-StA) has been cultured and is still capable of infecting plants (Flórez et al., 2017). The same strain produces antibacterial and antifungal compounds that can protect the beetle’s eggs from infection (Dose et al., 2018; Flórez et al., 2017). This is consistent with the hypothesis that the *B. gladioli* symbionts evolved from plant pathogens to become beetle mutualists. However, in field collections of *L. villosa*, Lv-StA is only found sporadically, and is never highly abundant (Flórez and Kaltenpoth, 2017). Instead, the most abundant strain is often the uncultured Lv-StB (Flórez and Kaltenpoth, 2017), which has been implicated in the production of the antifungal lagriamide, a defensive compound found in field egg collections (Flórez et al., 2018). We previously found through metagenomic sequencing that the genome of Lv-StB was much smaller than that of Lv-StA, suggesting it has undergone genome reduction (Flórez et al., 2018). It would seem that while *L. villosa* has multiple options for symbionts that produce potential chemical defenses, only a subset have specialized as obligate mutualists. The presence of multiple related strains in this system, with selective genome reduction of a single strain, could potentially shed light on why the genomes of some symbionts become reduced.

Here, we show that in the *L. villosa* microbiome, Lv-StB is uniquely undergoing genome reduction, despite other community members possessing biosynthetic pathways for potentially defensive molecules. We also suggest that this process was likely driven not only by horizontal acquisition of the putative lagriamide pathway, but also by loss of genes that limit cell division and translation, and gain of *zot*, a toxin also found in *Vibrio cholerae* that aids invasion of host membranes. Further, we present evidence that these horizontal gene transfers occurred concurrently with genome reduction, suggesting that complete genetic isolation is not a main driving force for the reduction process.

## RESULTS AND DISCUSSION

### Selective genome reduction of strain Lv-StB in *Lagria villosa*

We previously analyzed the metagenome of eight *L. villosa* egg clutches (Flórez et al., 2018), using our binning pipeline Autometa (Miller et al., 2019). This method has the advantage that it can separate noncharacterized eukaryotic contamination from metagenomes, and it uses multiple factors (nucleotide composition, sequence homology, the presence of single-copy marker genes and coverage) to accurately produce bins from individual datasets. Because we had implemented several bugfixes and small improvements to the pipeline since our original analysis, we re-ran Autometa on the same metagenomic assembly. Despite some minor differences, the new bins were broadly similar to our previous results (Dataset S1A), with 19 bins. As before, the Lv-StB bin had the highest coverage, at 1,977×, such that the constituent contigs are unlikely to be repeats from lower-coverage bins. We classified the bins according to a new standardized bacterial taxonomy utilized by the Genome Taxonomy Database (GTDB) that minimizes polyphyletic taxa and standardizes divergence between taxa of the same rank (Parks et al., 2018). Notably, the GTDB taxonomy reclassifies betaproteobacteria as being under class gammaproteobacteira. By this classification, all bins were in class Gammaproteobacteria, in three different orders: Betaproteobacteriales, Pseudomonadales and Xanthomonadales (Dataset S1B). The most abundant bins were all in the family Burkholderiaceae, with the highest abundance corresponding to the Lv-StB strain previously found to harbor the putative lagriamide biosynthetic gene cluster (BGC) (Flórez et al., 2018). Interestingly, the average nucleotide identity (ANI) of Lv-StB to the reference *B. gladioli* genome in GTDB (strain ATCC 10248) is 85.7%, much lower than the 95% cutoff suggested for species identifications (Goris et al., 2007) (Dataset S1B). This divergence suggests that Lv-StB is a novel species in the genus *Burkholderia*, even though we previously classified it as *B. gladioli* on the basis of 16S rRNA gene sequence (Flórez et al., 2018; Flórez and Kaltenpoth, 2017), and therefore we refer to the strain here as “*Burkholderia* Lv-StB”. Likewise, most bins were found to be novel species, with one (DBSCAN_round2_3) being divergent enough to be a representative of a novel genus in the family Burkholderiaceae. Notably, the cultured *B. gladioli* strain that we have previously isolated from *L. villosa* eggs, Lv-StA (Flórez et al., 2018, 2017), was not found to be present in this metagenome.

The bins obtained had a range of different sizes (Table 1), which could be due to either genome reduction or poor assembly and/or binning of a larger genome, which is often observed if there are many related strains in a metagenome (Miller et al., 2019). As part of the binning procedure, genome completeness was estimated based on the presence of 139 single-copy marker genes (Rinke et al., 2013) (Dataset S1C). However, as some complete genomes of genome-reduced symbionts have low apparent completeness by this measure (Miller et al., 2017, 2016b), this figure cannot be used alone to determine the size of an incompletely assembled genome. Conversely, even the drastically-reduced genomes of intracellular obligate insect symbionts have been found to almost universally maintain certain genes that we refer to here as ‘core genes’, involved in replication, transcription, protein folding/stability, tRNA modification, sulfur metabolism, RNA modification and translation (McCutcheon and Moran, 2012). We would expect, therefore, that a well-assembled reduced genome would contain a near complete core gene set, but not necessarily the whole set of 139 single-copy marker genes. Conversely, incompletely assembled genomes are likely to be missing a significant number of core genes that are required even in symbionts with highly reduced genomes. We examined the presence of core genes in all metagenomic bins, as well as *B. gladioli* Lv-StA for comparison (Dataset S1D and Table 1). Nine bins were close in size to the genome of their respective closest relative, while maintaining most core genes, and are classified as “nonreduced”, and ten bins were small but also lacked a significant fraction of core genes, and are classified as “incomplete”. Only the Lv-StB genome can be classified as reduced, on the basis of reduced size compared to its close relative (2.07 Mbp, 23.5%) and maintenance of most core genes (85.7%). This bin exhibited additional features of genomes undergoing reduction, namely reduced GC% compared to the *B. gladioli* reference genome (58.7% vs. 67.9%), and a proliferation of pseudogenes accounting for 45.29% of the annotated ORFs (Dataset S1C, S1E and Figure 1). Because of this, the Lv-StB genome exhibits a low coding density (59.04%), and it also possesses a large number of ORFs containing transposases (159). Both of these characteristics are hallmarks of symbionts in the early stages of genome reduction, where there are high numbers of pseudogenes and genome rearrangements (McCutcheon and Moran, 2012). The Lv-StB genome also contains a low number of genes compared with its free-living relative, with 744 ORFs that are not pseudogenes, transposases or hypothetical, versus 4,778 such genes in *B. gladioli* Lv-StA.

**Figure 1.**
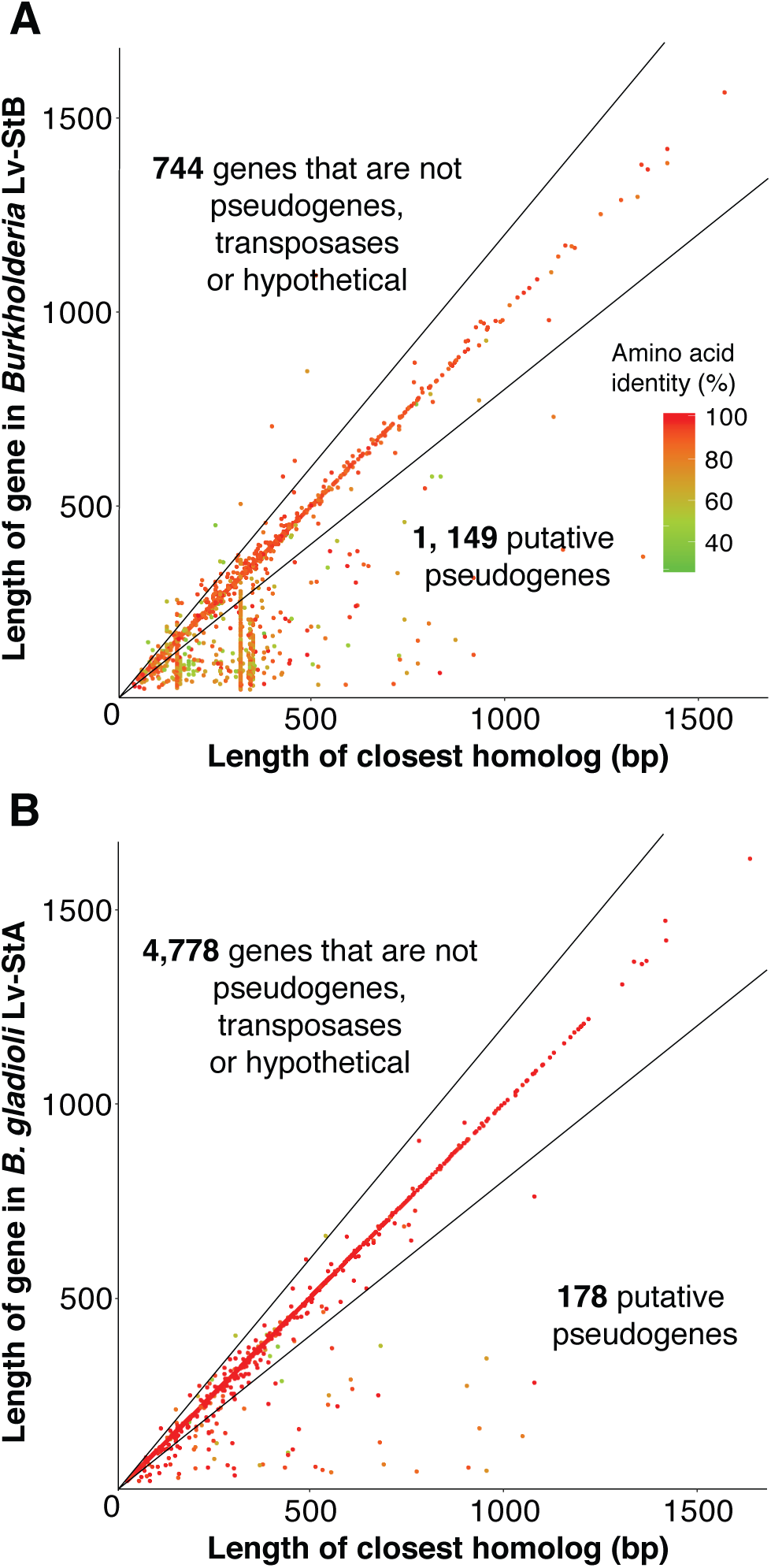
Comparison of the lengths of genes in the *Burkholderia* Lv-StB genome (**A**) and *B. gladioli* Lv-StA (**B**) with the closest homologs identified through BLAST searches against the NR database (see Methods). Genes which are less than 80% of the length of the closest relative (i.e. below the lower black line) are putatively assigned as pseudogenes, as described previously (Lerat and Ochman, 2005; Lopera et al., 2017). Note: Two vertical groupings of pseudogenes in Lv-StB correspond to multiple copies of a 155 bp hypothetical gene and an IS5 family transposase.

**Table 1.**
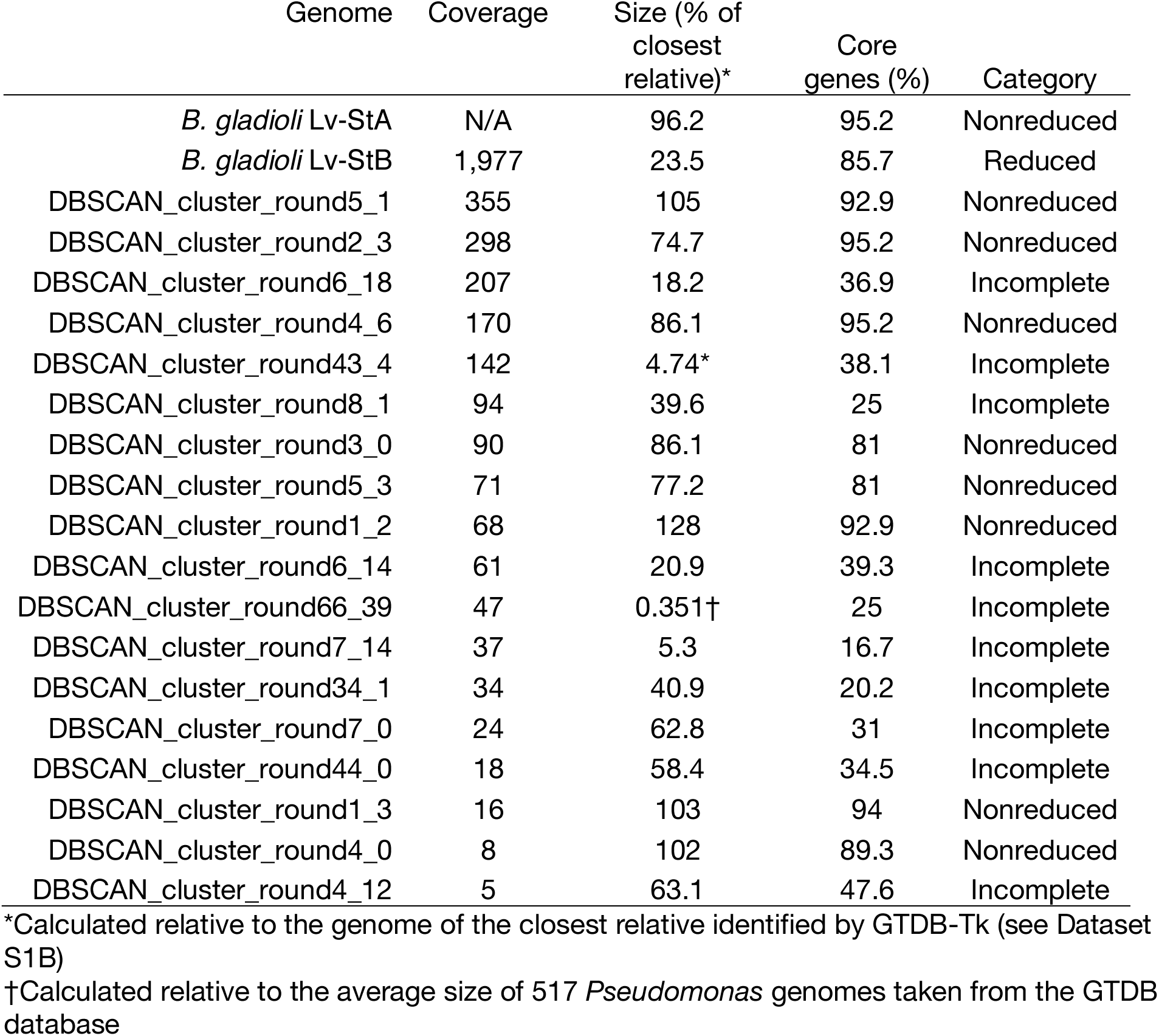
Genome characteristics of *Burkholderia* symbiont Lv-StB, its relative *B. gladioli* Lv-StA and other bins obtained from the metagenome.

| Genome | Coverage | Size (% of closest relative)* | Core genes (%) | Category |
| --- | --- | --- | --- | --- |
| *B. gladioli* Lv-StA | N/A | 96.2 | 95.2 | Nonreduced |
| *B. gladioli* Lv-StB | 1,977 | 23.5 | 85.7 | Reduced |
| DBSCAN_cluster_round5_1 | 355 | 105 | 92.9 | Nonreduced |
| DBSCAN_cluster_round2_3 | 298 | 74.7 | 95.2 | Nonreduced |
| DBSCAN_cluster_round6_18 | 207 | 18.2 | 36.9 | Incomplete |
| DBSCAN_cluster_round4_6 | 170 | 86.1 | 95.2 | Nonreduced |
| DBSCAN_cluster_round43_4 | 142 | 4.74* | 38.1 | Incomplete |
| DBSCAN_cluster_round8_1 | 94 | 39.6 | 25 | Incomplete |
| DBSCAN_cluster_round3_0 | 90 | 86.1 | 81 | Nonreduced |
| DBSCAN_cluster_round5_3 | 71 | 77.2 | 81 | Nonreduced |
| DBSCAN_cluster_round1_2 | 68 | 128 | 92.9 | Nonreduced |
| DBSCAN_cluster_round6_14 | 61 | 20.9 | 39.3 | Incomplete |
| DBSCAN_cluster_round66_39 | 47 | 0.351† | 25 | Incomplete |
| DBSCAN_cluster_round7_14 | 37 | 5.3 | 16.7 | Incomplete |
| DBSCAN_cluster_round34_1 | 34 | 40.9 | 20.2 | Incomplete |
| DBSCAN_cluster_round7_0 | 24 | 62.8 | 31 | Incomplete |
| DBSCAN_cluster_round44_0 | 18 | 58.4 | 34.5 | Incomplete |
| DBSCAN_cluster_round1_3 | 16 | 103 | 94 | Nonreduced |
| DBSCAN_cluster_round4_0 | 8 | 102 | 89.3 | Nonreduced |
| DBSCAN_cluster_round4_12 | 5 | 63.1 | 47.6 | Incomplete |

\*Calculated relative to the genome of the closest relative identified by GTDB-Tk (see Dataset S1B)
†Calculated relative to the average size of 517 *Pseudomonas* genomes taken from the GTDB database

### Diversity of biosynthetic gene clusters in the *L. villosa* microbiome

Because we had previously isolated the non-reduced *B. gladioli* Lv-StA strain from *L. villosa* (Flórez et al., 2018), and found it to produce protective compounds despite having sporadic distribution and low abundance in field-collected beetles, we asked whether BGCs were a unique feature of Lv-StB in the metagenome, or whether other community members have the biosynthetic machinery for potential chemical defenses. AntiSMASH (Blin et al., 2017) searches revealed a total of 105 BGCs in the metagenome (Fig. 2), with variable BGC content in the bins, from zero to 566 kbp (0 to 14 BGCs, with 16 BGCs in the unclustered bin), while the *B. gladioli* Lv-StA genome contained 1006 kbp BGCs (21 BGCs). This indicates that Lv-StB is not the only strain in the egg microbiome with the potential to produce complex natural products, and many strains harbor multiple BGCs. In the metagenome, bins in the family Burkholderiaceae collectively contained the most BGCs by length (963 kbp), followed by a single bin in family Rhodanobacteraceae (DBSCAN_round1_2, 566 kbp, Fig. 2). This distribution suggests that although Burkholderiaceae appear to be an important reservoir of BGCs in the *L. villosa* egg microbiome, other groups have significant biosynthetic potential. Out of the 126 BGCs detected in the metagenome and in the *B. gladioli* Lv-StA genome, there were 17 > 50 kbp in length predicted to produce complex nonribosomal peptides or polyketides (Fig. 3). Two of these have been putatively assigned to production of the antibiotic lagriene in Lv-StA (Flórez et al., 2017), and the antifungal lagriamide in Lv-StB (Flórez et al., 2018), whereas five other small molecules known to be produced by Lv-StA have been assigned to shorter BGCs (Dose et al., 2018; Flórez et al., 2017). That 15 out of the 17 largest assembled BGCs remain without characterized products suggests that the majority of biosynthetic pathways in the *L. villosa* egg microbiome likely codes for novel small molecule products. We compared the 126 identified BGCs using BIG-SCAPE (Navarro-Muñoz et al., 2018), and found only 7 examples of BGCs occurring in multiple strains, indicating that the biosynthetic potential in the metagenome and *B. gladioli* Lv-StA is largely nonredundant. Taken together, this suggests that there is a large amount of undefined biosynthetic potential for small molecule production in *L. villosa* symbionts, beyond *B. gladioli* Lv-StA and *Burkholderia* Lv-StB.

**Figure 2.**
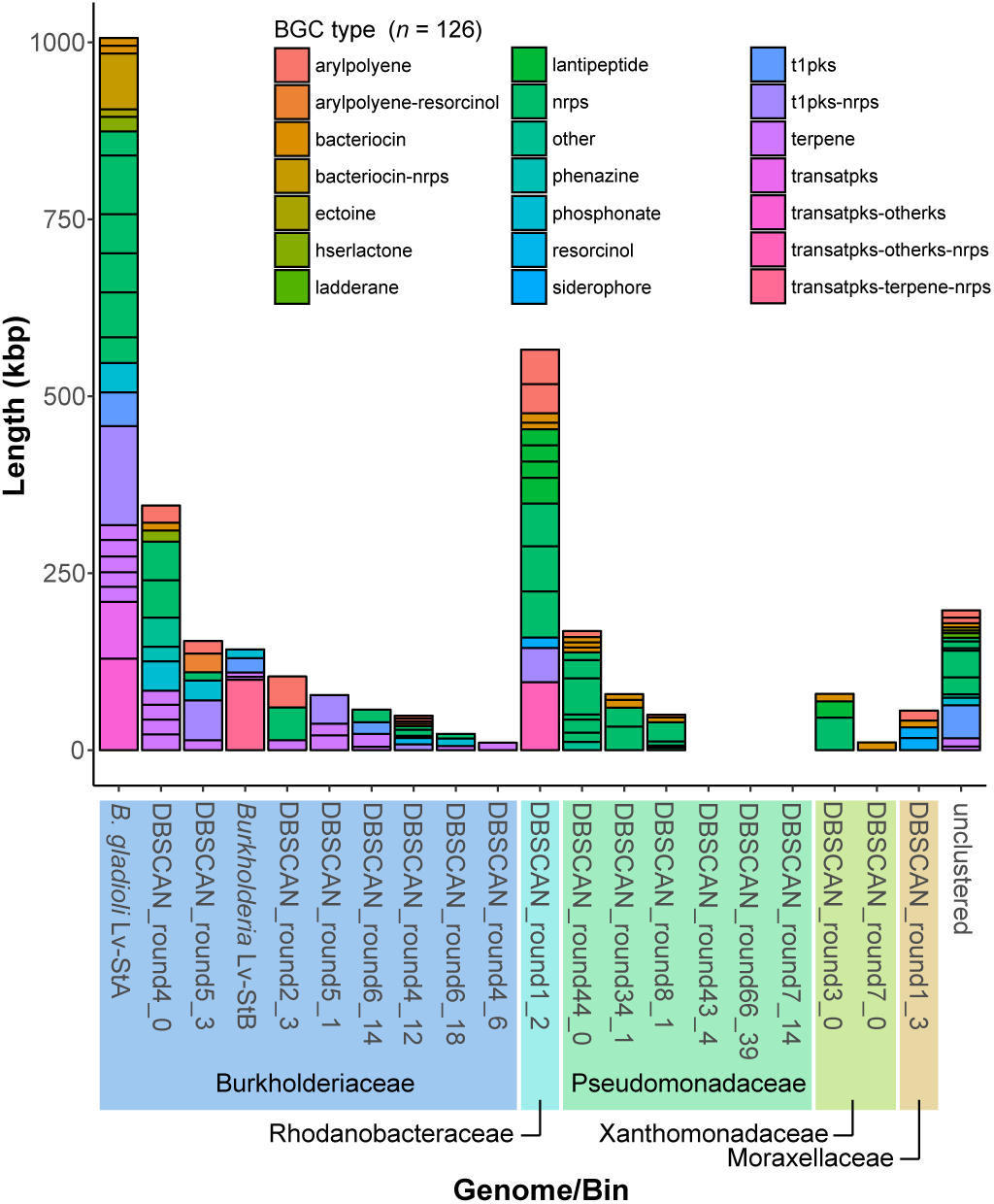
Distribution of biosynthetic gene clusters (BGCs) amongst the *L. villosa* metagenome bins and the genome of *B. gladioli* Lv-StA. Colors indicate the type of BGC annotated by antiSMASH (126 identified) (Blin et al., 2017).

**Figure 3.**
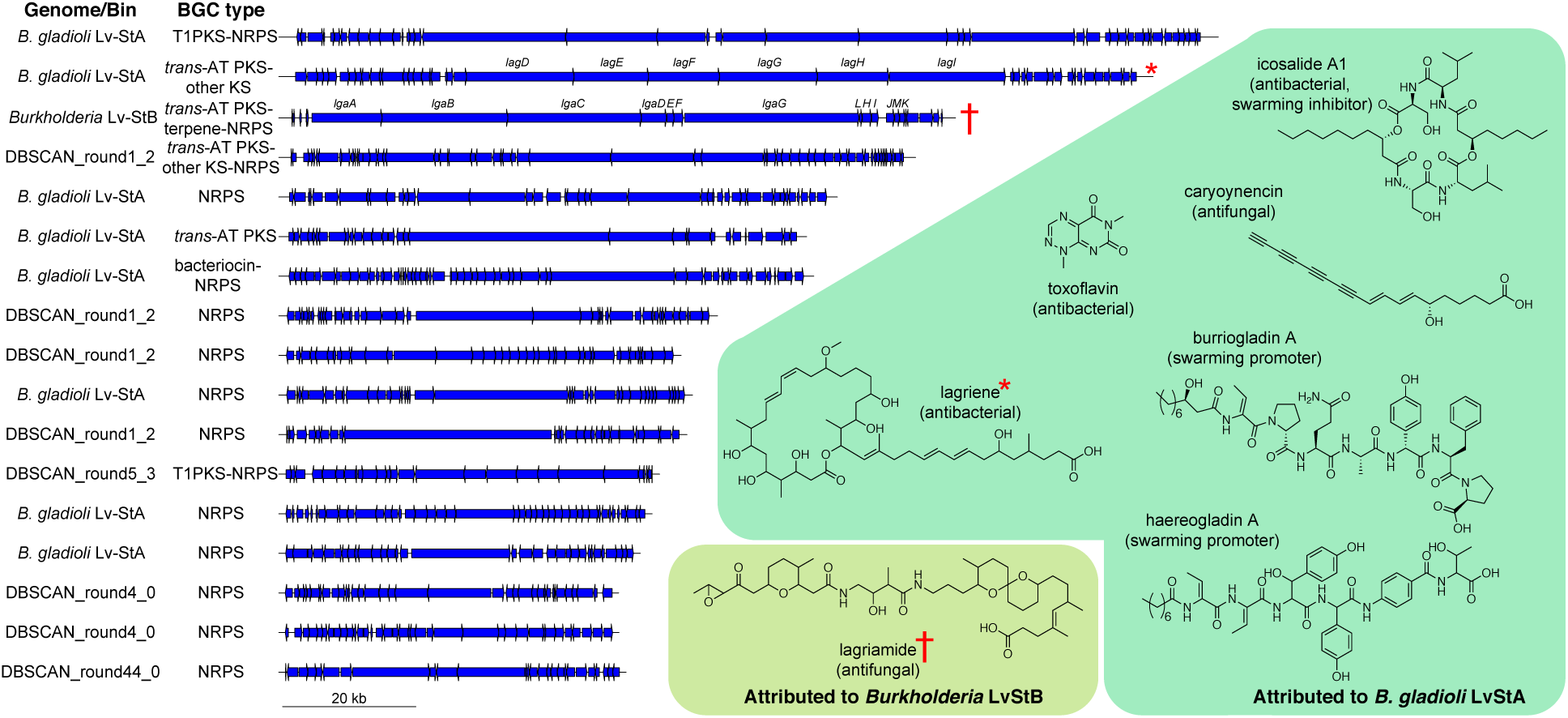
Schematics of all BGCs with greater than 50 kbp length assembled from the *L. villosa* metagenome and in the *B. gladioli* Lv-StA isolate genome, out of 126 identified by antiSMASH (left). Shown on the right are structures of compounds putatively assigned to BGCs in the *B. gladioli* Lv-StA genome (Dose et al., 2018; Flórez et al., 2017), or putatively assigned to a BGC in the *Burkholderia* Lv-StB genome (Flórez et al., 2018). Structures highlighted with red symbols have been attributed to the indicated BGCs.

### Divergence between Lv-StB and the closest free-living relative

We constructed a phylogenetic tree of metagenomic bins assigned to the genus *Burkholderia* as well as *B. gladioli* Lv-StA, based on 120 marker genes (Fig. 4A). This showed that *Burkholderia* Lv-StB is most highly related to the *B. gladioli* clade but is divergent from it. We calculated genome-wide ANI values for pairs of *Burkholderia* genomes, and found *B. gladioli* strains shared between 97-100% ANI, while at most *Burkholderia* Lv-StB shared 85.79% ANI with *B. gladioli* A1 (Dataset S1F). During genome reduction, symbionts are known to undergo rapid evolution due to the loss of DNA repair pathways (as found in the Lv-StB genome, see below) and the relaxation of selection (McCutcheon and Moran, 2012), and so the divergence of Lv-StB from *B. gladioli* may have been accelerated relative to free-living lineages. Genome-reduced symbionts have often been vertically transmitted for evolutionary timescales and across host speciation events, and therefore it is possible to calculate evolution rates where related symbionts occur in hosts with known divergence times inferred from fossil records. Such estimates in insect symbionts vary widely over three orders of magnitude, but more recent ant and sharp-shooter symbiont lineages (established < 50 Mya for “*Candidatus* Baumannia cicadellinicola”, BAU; *Blochmannia obliquus*, BOB; *Bl. pennsylvanicus*, BPN and *Bl. floridanus*, BFL) show high rates of divergence per synonymous site per year (dS/t) between 1.1 × 10^-8^ and 8.9 × 10^-8^ (Silva and Santos-Garcia, 2015) (the divergence rates used here are found in Table S1). Because of the large number of pseudogenes in the Lv-StB genome, we reasoned that it is likely to be a recent symbiont, and therefore used these rates to estimate divergence times between Lv-StB and *B. gladioli* A1. We found a dS of 0.5486 per site between these genomes, and calculated divergence times of 6.15, 8.55, 6.93 and 49.76 My for rates BFL, BPN, BOB and BAU, respectively (Table S1). We should note here that these figures are very approximate, and are possibly overestimates as symbiont evolution rates are likely not constant, with particularly rapid evolution occurring during lifestyle transitions (Lo et al., 2016). The range of these estimated divergence times suggests that the common ancestor of *Burkholderia* Lv-StB and *B. gladioli* existed after the evolution of symbiont bearing structures in Lagriinae beetles (see below).

**Figure 4.**
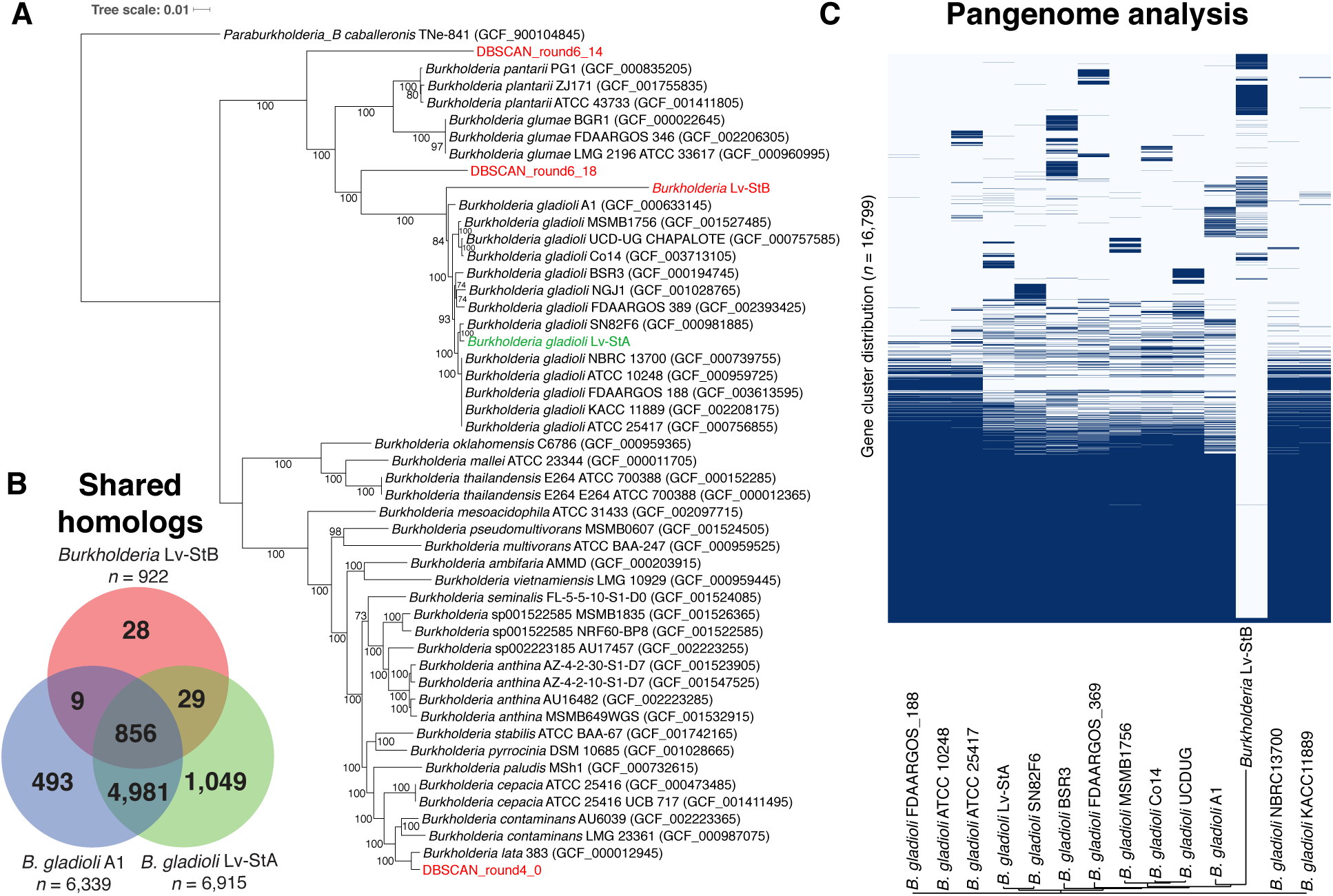
(**A**) Maximum-likelihood multilocus species tree of metagenomic bins classified in the genus *Burkholderia*, plus *B. gladioli* Lv-StB, using 120 marker gene protein sequences. Bootstrap proportions greater than 70% are expressed to the left of each node as a percentage of 1,000 replicates. Metagenomic bins are shown in red, while the *L. villosa*-associated isolate *B. gladioli* Lv-StA is shown in green. (**B**) Shared homologous gene groups in *Burkholderia* Lv-StA, *B. gladioli* A1 and *B. gladioli* Lv-StB, after discounting pseudogenes. Note: Homologous groups are counted only once per genome (i.e. collapsing paralogs), and therefore counts are lower than absolute gene counts. Also, 492 genes in the Lv-StB genome were singletons and not included in homologous groups. (**C**) Hierarchical clustering of homologous gene clusters, showing presence and absence in *Burkholderia* Lv-StB and closely-related strains of *B. gladioli*.

We then sought to quantify the conservation of genes in Lv-StB compared to 13 related *B. gladioli* strains, by identifying homologous gene groups among the entire set of non-pseudogenes in these strains with OMA (Altenhoff et al., 2018). This pipeline aims to identify orthologous groups while discounting paralogs among the genes in a given set of genomes (Dessimoz et al., 2005). Of the 1,388 genes in Lv-StB that are not pseudogenes (Dataset S1C), 492 were not included in any OMA orthologous group (see below). A crude analysis of the OMA groups in Lv-StB, *B. gladioli* Lv-StA and *B. gladioli* A1 revealed that Lv-StB retains a small subset of groups found in both Lv-StA and A1, and has few unique groups (Fig. 4B), suggesting that Lv-StB has lost many of the genes conserved in *B. gladioli*. Consistent with this notion, we visualized the pangenome of *B. gladioli* and Lv-StB with Roary (Page et al., 2015) (Fig. 4C) and found a large number of gene clusters that are conserved in *B. gladioli*, but not Lv-StB. The gene clusters that are more variable amongst *B. gladioli* are also generally not found in Lv-StB. Conversely, there were gene clusters found in Lv-StB that are not present in *B. gladioli* strains, and these clusters may have been obtained by horizontal transfer after the divergence of Lv-StB, or alternatively were lost in *B. gladioli*.

Remarkably, out of the 1,149 pseudogenes detected in the Lv-StB genome, 976 were hypothetical and 129 were transposases, leaving only 44 that were recognizable (Dataset S1G). This set of pseudogenes included a number of important genes in the categories noted to be depleted below. For instance, the DNA polymerase I gene (*polA*) appears to have been disrupted by a transposase, which is now flanked by two DNA pol I fragments (E5299_1120 and E5299_01122). Likewise, the *uvrC* gene (E5299_00503), a component of the nucleotide excision repair system (Lin and Sancar, 1992), is also present as a truncated gene adjacent to a transposase. There were also pseudogenes involved in the Entner-Doudoroff and glycolysis energy-producing pathways (phosphogluconate dehydratase (Carter et al., 1993) and glucokinase (Lunin et al., 2004)), as well as purine biosynthesis (phosphoribosylglycinamide formyltransferase (Almassy et al., 1992), phosphoribosylformylglycinamidine synthase (Schendel et al., 1989)). Interestingly, we found two pseudogenes that negatively regulate cell division and translation. Septum protein Maf is a nucleotide pyrophosphatase that has been shown to arrest cell division, especially after transformation or DNA damage (Tchigvintsev et al., 2013). Deletion of the *E. coli* gene for homolog YhdE increased growth rate, while overexpression decreased growth rate (Jin et al., 2015). Therefore the loss of Maf in Lv-StB would be expected to increase the rate of cell division and reduce the conversion of nucleotides, which it probably obtains from the host, to the monophosphates. The gene for the energy-dependent translational throttle protein EttA was also found to be truncated. This protein slows translation through interacting with the ribosome in both the ATP- and ADP-bound forms (Boël et al., 2014; Chen et al., 2014). Under energy-depleted conditions (i.e. high ADP), EttA was found to stabilize ribosomes and prevent commitment of metabolic resources, and thus the deletion mutant displayed reduced fitness during extended stationary phase (Boël et al., 2014). However, under circumstances where the host supplies ample nucleotides, the loss of *ettA* would be expected to increase translation rates.

### Degradation of primary metabolic pathways in *Burkholderia* Lv-StB

Lv-StB is deficient in many metabolic pathways that are complete in related *B. gladioli* strains (Fig. 5), including the glyoxylate shunt (Dolan and Welch, 2018), various carbon degradation pathways, mixed acid fermentation, as well as sulfur and nitrogen metabolism. Although the extent of gene losses could be overestimated due to the draft status of the Lv-StB genome, the pervasiveness of metabolic gaps found combined with the high coverage of the genome (Dataset S1A) suggest generalized gene loss in many functional categories. Lv-StB appears incapable of making any of the following compounds due to the absence of several biosynthetic genes: Thiamine, riboflavin, nicotinate, pantothenate, vitamin B12 and biotin. Likewise, there were deficiencies in amino acid biosynthesis (Fig. S1). We predict that Lv-StB would be able to make chorismate, isoleucine, leucine, ornithine, proline and threonine, but likely lacks the ability to make aromatic amino acids, serine, methionine, lysine, histidine, cysteine, glutamine and arginine due to the absence of several key genes in these pathways. The genome of Lv-StB also lacks genes involved in chemotaxis and flagella, suggesting that after the symbiont mixture is spread on eggs, the colonization of the dorsal cuticular structures in the embryo (Flórez et al., 2017) does not require symbiont motility. Interestingly, the Lv-StB genome includes a trimeric autotransporter adhesin (TAA) related to SadA (Raghunathan et al., 2011), which is involved in the pathogenicity of *Salmonella typhimurium* by aiding cell aggregation, biofilm formation, and adhesion to human intestinal epithelial cells. TAAs are found in Proteobacteria and consist of anchor, stalk and head domains, of which the head forms the adhesive component (Linke et al., 2006). The bacterial honey-bee symbiont, *Snodgrassella alvi* is hypothesized to utilize TAAs in combination with other extracellular components during colonization of the host gut (Powell et al., 2016), and similar genes were identified in *S. alvi* symbionts in bumble bees and are predicted to perform a similar role (Kwong et al., 2014). Therefore, this gene may play a role in the adhesion of Lv-StB cells to *L. villosa* eggs.

**Figure 5.**
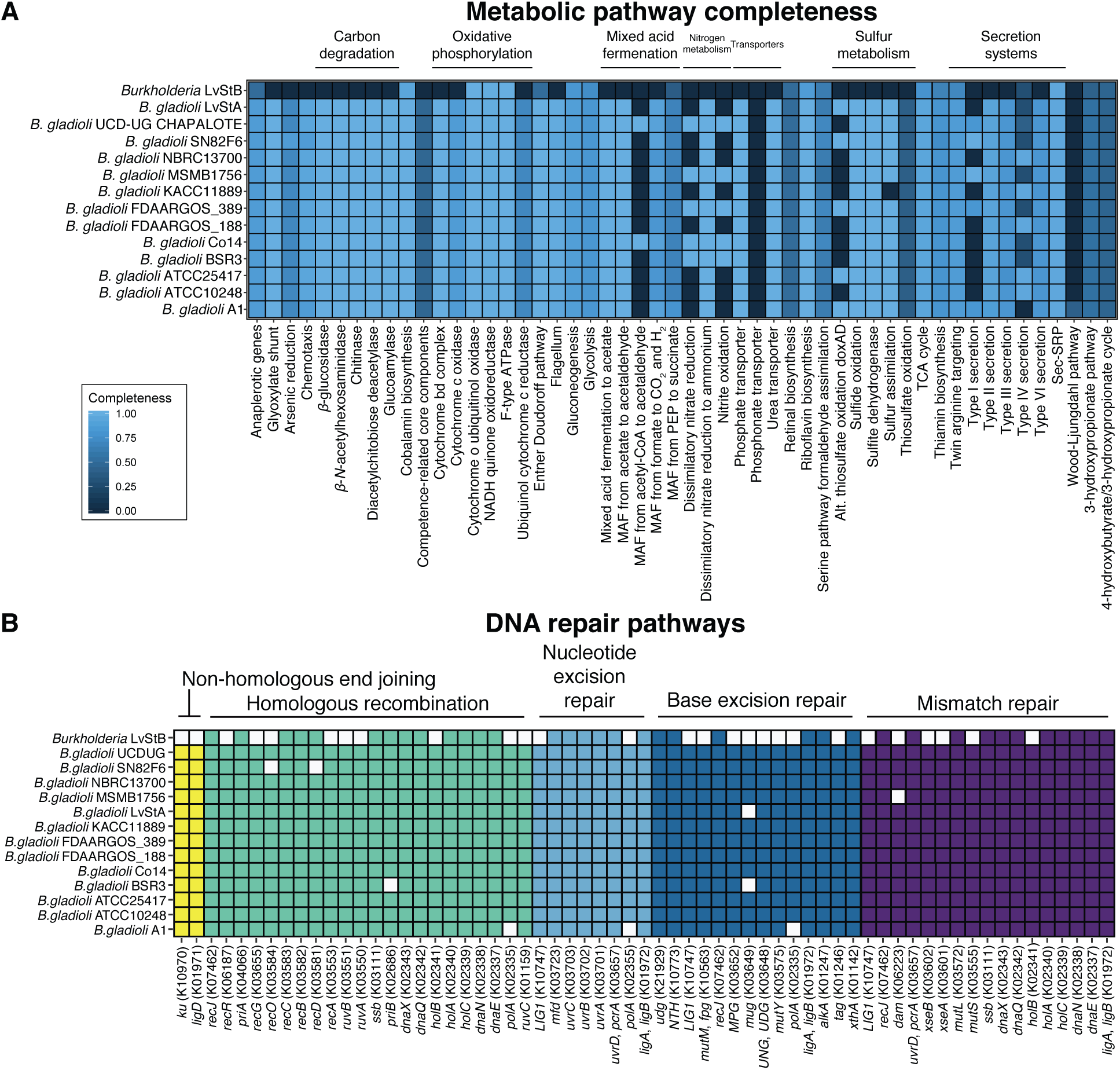
Completeness of metabolic and DNA repair pathways in *Burkholderia* Lv-StB in comparison to closely-related strains of *B. gladioli*. (**A**) Completeness of various metabolic pathways as determined by KEGG-decoder. Note: Categories that were not found in any of the examined genomes have been removed. (**B**) The presence (colored squares) and absence (white squares) of genes in the different DNA repair pathways in Lv-StB and related *B. gladioli* strains.

The Lv-StB genome is also missing several enzymes within glycolysis, most notably phosphoglycerate kinase, phosphoenolpyruvate carboxykinase and others, which would suggest that Lv-StB has lost the ability to perform glycolysis. The loss of glycolysis is often substituted by an alternative pathway, such as the pentose phosphate or Entner-Doudoroff pathway (Chen et al., 2016). This is not the case for Lv-StB, where both the glucose-6-phosphate dehydrogenase and 6-phosphogluconolactonase genes appear to be missing from the genome. The citrate cycle is largely complete, except that it is missing pyruvate carboxylase, the enzyme that converts pyruvate to oxaloacetate. However, phosphoenolpyruvate carboxylase is present in Lv-StB (E5299_00983) and may alternatively be used for the production of oxaloacetate from exogenous phosphoenol-pyruvate (Takeya et al., 2017) as alternative pathways for the supply of oxaloacetate are also incomplete. Lv-StB is also missing all genes required to form cytochrome c oxidase and cytochrome b-c complexes. However, all genes required to encode NADH:quinone oxidoreductase (Complex I), succinate dehydrogenase (Complex II) and cytochrome o ubiquinol oxidase are present, along with all genes required for the F-type ATPase. The lack of Complex III would likely result in a decreased rate of ATP production in Lv-StB as observed in fungi with alternative oxidases that bypass Complexes III and IV (Duarte and Videira, 2009). Lv-StB may be similar to the psyllid endosymbiont “*Candidatus* Liberibacter asiaticus”, which has lost both key glycolysis and glyoxalase genes and instead relies on the scavenging of ATP from the host (Jain et al., 2017).

We also found many deficiencies in both the *de novo* and salvage nucleotide pathways (Fig. S2). In the pyrimidine biosynthetic pathway, genes for dihydrooratase (*pyrC*) (Porter et al., 2004) and orotate phosphoribosyl transferase (*pyrE*) (Aghajari et al., 1994) were missing, suggesting that Lv-StB cannot produce orotate from *N-* carbamoylaspartate, and cannot create nucleotides from free pyrimidine bases. Ribonucleotide reductase (*nrdAB*) (Brignole et al., 2012) and thymidylate synthase (*thyA*) (Carreras and Santi, 1995) are present, suggesting that deoxypyrimidine nucleotides can be made from CTP. The deficiencies in purine synthesis were more profound. The majority of the *de novo* pathway (Nelson et al., 2008) was missing (*purCDEFHLMNT*, IMP dehydrogenase/*guaB*), except for adenylosuccinate lyase (*purB*), adenylosuccinate synthase (*purA*) and GMP synthase (*guaA*). Lv-StB should therefore be able to make AMP from IMP, and GMP from XMP (plus their deoxy analogs through ribonucleotide reductase), but cannot make purines *de novo*. We were also not able to find adenine phosphoribosyltransferase or hypoxanthine-guanine phosphoribosyltransferase (Nelson et al., 2008), meaning that purine bases cannot be salvaged to make nucleotides.

### Degradation of DNA repair pathways in *Burkholderia* Lv-StB

The genome of Lv-StB is missing many genes involved in DNA repair (Fig. 5B), similar to other examples of genome-reduced symbionts (McCutcheon and Moran, 2012). Compared to closely-related *B. gladioli* strains, Lv-StB lacks genes in every repair pathway. In particular DNA polymerase I (*polA*), used in homologous recombination, nucleotide excision repair and base excision repair, is only present as two truncated pseudogenes (see above). Even though *polA* is involved in many different DNA repair pathways, it has been found to be nonessential in *Escherichia coli* (Gerdes et al., 2003; Goodall et al., 2018), *B. pseudomallei* (Moule et al., 2014) and *B. cenocepacia* (Higgins et al., 2017). In the homologous recombination pathway, Lv-StB lacks *recA*, *polA*, *ruvA*, *ruvB*, *ruvC* and *recG*, all of which have been found to be essential for homologous recombination in *E. coli* (Kowalczykowski et al., 1994). Likewise, Lv-StB is also missing the two components of the nonhomologous end-joining pathway, *ku* and *ligD* (Pitcher et al., 2007), suggesting that it cannot recover from double-strand breaks. In the base excision repair pathway, Lv-StB lacks several DNA glycosylases which are responsible for removing chemically modified bases from double-stranded DNA (McCullough et al., 1999). Some of these losses simply reduce redundancy, but it has also lost nonredundant glycosylases *mutM* and *mug*. The former recognizes 2,6-diamino-4-hydroxy-5-*N*-methylformamidopyrimidine (Fapy) and 8-hydroxyguanine (Boiteux et al., 1992), while the latter recognizes G:U and G:T mismatches (Barrett et al., 1998) as well as epsilonC (Saparbaev and Laval, 1998), 8-HM-epsilonC (Hang et al., 2002), 1,N(2)-epsilonG (Saparbaev et al., 2002) and 5-formyluracil (Liu et al., 2003). Finally, in the mismatch repair system, Lv-StB is missing *mutS*, which is required for the recognition of mismatches in methyl-directed repair (Schofield and Hsieh, 2003). In summary, Lv-StB is likely to be completely incapable of nonhomologous end-joining, homologous recombination, and mismatch repair, while being impaired in nucleotide excision repair and base excision repair due to the loss of DNA polymerase I and several DNA glycosylases.

### Timing of horizontal acquisition of defensive and other genes in the Lv-StB genome

We then attempted to identify genes in the *Burkholderia* Lv-StB genome that are likely to have been acquired by horizontal gene transfer (HGT). A total of 497 non-pseudogenes were identified as unique to Lv-StB, and following removal of genes with no matches against the BLAST NR database, and removal of genes that were homologous to other *B. gladioli* genomes not included in this study, there were 148 genes that appeared to be more closely related to species other than *B. gladioli*, that have potentially been acquired through horizontal transfer (Figure 6A). Most genes appear to have been obtained from gammaproteobacteria and alphaproteobacteria, with a small number from firmicutes, cyanobacteria and phages (see Dataset S1H). The distribution is consistent with the notion that horizontal transfer occurs most frequently between closely-related species (Gillings, 2017). In particular, Burkholderiaceae was the most frequent apparent donor amongst gammaproteobacterial proteins. Interestingly, the alphaproteobacterial genus *Ochrobactrum* (family Brucellaceae in the NCBI taxonomy, Rhizobiaceae in GTDB) was a major putative gene donor. This genus includes several symbionts of termites (Mathew et al., 2012), army worms (Jones et al., 2019), weevils (Montagna et al., 2015) and leeches (McClure et al., 2019; Rio et al., 2009). In previous 16S amplicon investigations of *Lagria* beetles, *Ochrobactrum* were often found (Flórez et al., 2017; Flórez and Kaltenpoth, 2017) (Fig. S3), suggesting that the donors of these genes could have also been associates of *L. villosa*. *Ochrobactrum* OTUs account for 5–20% of 16S rRNA gene reads, but this genus was not observed in the shotgun metagenome. However, disparities between 16S and shotgun metagenome abundances are not uncommon due to variable 16S copy number, primer and sequencing biases (Chen et al., 2017; Delforno et al., 2017). Based on the evidence for putatively horizontally transferred genes, we asked whether these could have contributed to the dominance of Lv-StB in *L. villosa*, and set out to estimate the timings of horizontal transfer events.

**Figure 6.**
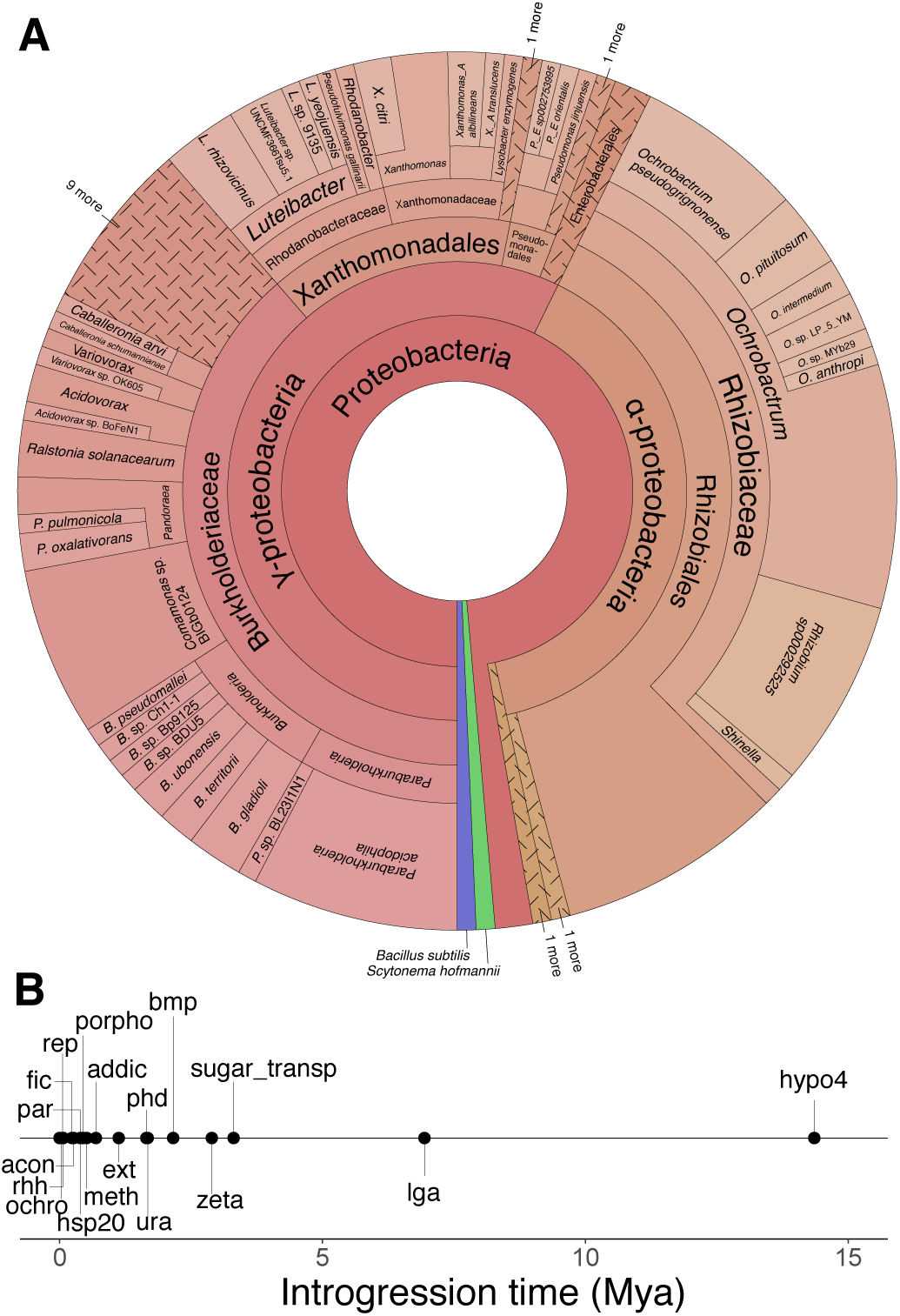
**(A)** Putative sources of genes unique to the *Burkholderia* Lv-StB genome (compared to *B. gladioli* strains) based on hits to BLASTP searches. Note: Three proteins of putative phage origin (see Dataset S1I) are not included in the figure. **(B)** Estimated introgression time for putative HGT gene sets, using the “BAU” substitution rates (Silva and Santos-Garcia, 2015). Note: For clarity, the gene sets with estimated introgression time of < 5,000 ya are not labeled. For these and ages estimated with other substitution rates, see Dataset S1I.

Horizontally transferred genes are often detected on the basis of nucleotide composition differing from other genes in the genome (Becq et al., 2010). Such genes initially exhibit nucleotide composition consistent with the donor genome, which over time will eventually normalize to the composition of the recipient genome (Lawrence and Ochman, 1997). The rate of this “amelioration” process (ΔGC^HT^) has been modeled by Lawrence and Ochman (Lawrence and Ochman, 1997), based on the substitution rate (*S*), the transition/transversion ratio (κ) and GC content of both the recipient genome (GC^EQ^) and putatively horizontally transferred genes (GC^HT^), according to equation 1. By iterating this equation repeatedly until GC^HT^ equals GC^EQ^, the time required from the present day to complete amelioration can be estimated. If the GC content of the donor genome is known, then equation 1 can be used in reverse to estimate the time since introgression. However, if the donor GC content is not known, then the differing selection pressures on the first, second and third codon positions can be exploited to estimate the introgression time. Because these positions have different degrees of amino acid degeneracy, they are subject to different degrees of selection, and therefore they ameliorate at different rates. As a consequence, Lawrence and Ochman (Lawrence and Ochman, 1997) found that for genes in the process of amelioration, the relationship between overall GC content and the GC content at individual codon positions seen in genes at equilibrium (Lawrence and Ochman, 1997; Muto and Osawa, 1987) (Equations 2, 3 and 4) does not hold. So if equation 1 is applied in reverse separately for each codon position, the time since introgression can be inferred at the iteration yielding the minimum square difference from equations 2–4. Application of equation 1 also yields an estimate for the original donor GC.

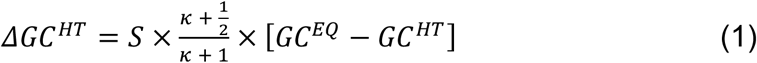

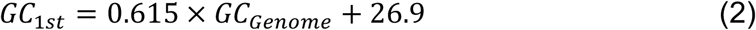

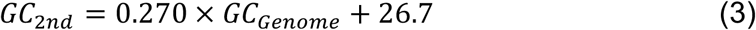

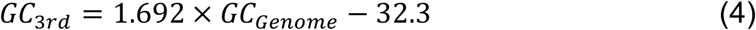

We identified groups of consecutive genes in the putative HGT set that could have been acquired together, and used the above method to estimate their introgression time (Dataset S1H). Out of the 18 identified gene groups, 7 were found to have atypical GC content (defined by Lawrence and Ochman (Lawrence and Ochman, 1997) as either >10% lower or >8% higher GC% in first or third codon positions compared to the genome as a whole). The method above was used to estimate time of introgression for each gene group, using the BFL, BPN, BOB and BAU divergence rates (see above, Fig. 6B, Dataset S1H). The oldest horizontal transfer was found to be group “hypo4” (1.79–14.36 Mya), representing two phage proteins, and the next oldest is the lagriamide BGC and neighboring genes (*lga*, 0.8–6.42 Mya). The *lga* BGC is predicted to have come from a high-GC organism, with an original GC content of 72%. The closest relatives of many of the *lga* genes are found in *Pseudomonas* strains, which typically do not have GC contents this high. However, as BGCs are often thought to be horizontally transferred (Jensen, 2016), *Pseudomonas* may not be the direct source of *lga* in Lv-StB.

Several other HGT gene groups were predicted to be involved in transport (sugar_transp, bmp, ext, dinj, tonb, tonb2), with predicted substrates including purine nucleosides, trehalose and vitamin B12. Of these, the sugar_transp and bmp groups predicted to be involved in trehalose and purine import, respectively, are relatively old (0.41–3.31 My and 0.27–2.16 My), and the groups likely involved in B12 import (dinj, tonb, tonb2) are estimated to be <5,000 y old. Trehalose is the most abundant component of insect hemolymph, and in a leaf beetle system was found to be provisioned by the host to its genome-reduced symbiont (Bauer et al., 2019). We also found HGT gene groups putatively involved in heat shock response (hsp20, 0.05–0.4 Mya), and DNA repair (ura, rep, ochro). In the latter category was a uracil-DNA glycosylase (in ura, 0.21–1.68 Mya) used in base excision repair in case of deamination, RecB used in homologous recombination (in rep, 0.01–0.055 Mya) (Nelson et al., 2008) and YedK (in ochro, 0.01-0.035 Mya), a protein used in the SOS response that binds to abasic sites in single strand DNA (Mohni et al., 2019). The transferred transport functions and DNA repair proteins match functional categories that are currently lacking genes due to genome reduction (see above), and therefore the transfers could have been contemporaneous with the reduction process, acting as compensatory mechanisms for lost functions.

One putative HGT toxin that is unique to Lv-StB in the metagenome and amongst *B. gladioli* strains is zonular occludens toxin (zot, estimated as acquired <5,000 ya). The *zot* gene is responsible for the production of the zonula occuludens toxin, a virulence factor which was initially identified in *Vibrio cholera* and was found to lead to the disassembly of intracellular tight junctions, leading to increased permeability of mammalian epithelium (Di Pierro et al., 2001). Co-localized with the predicted zot gene was a gene encoding DUF2523, which we found often accompanied *zot* in searches of the STRING database (Szklarczyk et al., 2017). Zot proteins have been identified in several strains of *Camplylobacter* and have been shown to elicit an inflammatory response in intestinal epithelial cells (Liu et al., 2016; Mahendran et al., 2016). Furthermore, a significant correlation was found between the presence of the Zot protein and hyper-invasive strains of *Neisseria meningitides* (Joseph et al., 2011). Potentially, *zot* may aid in the infection of the *L. villosa* embryonic structures through increasing permeability across the outer layers of the egg, although this remains speculative.

## CONCLUSIONS

In the 1920s it was observed (Jürgen Stammer, 1929) that beetles in the *Lagria* and *Cerogria* genera contained structures now known to harbor *Burkholderia* symbionts in *L. villosa* and *L. hirta* (Flórez and Kaltenpoth, 2017). Other genera in the Lagriinae subfamily, such as *Adynata* and *Arthromacra*, do not contain these structures. According to the tree found at timetree.org (Kumar et al., 2017), *Lagria* and *Cerogria* diverged 55 Mya, and the common ancestor of *Lagria*, *Cerogria* and *Adynata* existed 82 Mya (this region of the tree utilizes data from Kergoat *et al*. (Kergoat et al., 2014)). Based on these estimates, the symbiont-bearing structures in *Lagria* and *Cerogria* likely evolved between 82 and 55 Mya. Our analysis suggests that the divergence of *Burkholderia* Lv-StB from *B. gladioli* occurred after that point (6.15–49.76 Mya). During that time, the genome of *Burkholderia* Lv-StB became reduced, and it is likely dependent on the host due to deficiencies in energy metabolism and nucleotide biosynthesis. Notably, the profound metabolic insufficiencies and incomplete DNA repair pathways in Lv-StB are typical of symbionts with smaller genomes, such as “*Ca.* Endolissoclinum faulkneri”, an intracellular tunicate symbiont with a 1.48 Mbp genome and a similar number of genes (783) (Kwan et al., 2012), estimated to have been a symbiont for at least 6-31 My (Kwan and Schmidt, 2013). While the presence of *Burkholderia* Lv-StB and its defensive compound lagriamide has been shown to decrease the rate of fungal egg infection (Flórez et al., 2018), the symbiont is not essential for beetle reproduction (Flórez et al., 2017). Therefore, the relationship is facultative from the perspective of the host, while Lv-StB is in the process of becoming dependent on *L. villosa*. A central question we aimed to answer in this work was how the genome of Lv-StB became reduced, when *L. villosa* appears to maintain multiple other nonreduced *Burkholderia* and other symbionts.

It is clear from previous work that the *Lagria* symbionts related to *B. gladioli* evolved from plant associated strains (Flórez and Kaltenpoth, 2017), likely transmitted to the insects from the plant environment. The probable advantage of this early association for the hosts was protection of eggs from infection, through small molecules made by its microbiome. The strains characterized here, as well as the previously isolated Lv-StA, were found to contain ample biosynthetic potential, and both Lv-StA and Lv-StB produce antifungal compounds that protect eggs from fungal infection in lab experiments (Flórez and Kaltenpoth, 2017). Yet Lv-StA is only found sporadically in the field as a minor component of the microbiome (Flórez et al., 2018). It is probably advantageous for *Lagria* beetles to maintain a pool of facultative symbionts with different biosynthetic capability, to allow for fast adaptation to different environmental infection pressures (Flórez et al., 2015). However, there may be less selection pressure on a facultative symbiont to stay associated with its host if it can also survive in the environment and infect plants.

The foundational event in the establishment of the symbiosis between Lv-StB and *L. villosa* was likely the acquisition of the *lga* pathway, which putatively produces lagriamide, in a non-reduced ancestor of Lv-StB (Flórez et al., 2018). We place this as the first event for four reasons. First, the *lga* BGC is almost the oldest detectable horizontal transfer that survives in the reduced genome of Lv-StB. Second, we found little evidence that Lv-StB is capable of making metabolites of use to the host, indicating that the symbiosis is likely not based on nutrition. *L. villosa*’s diet of plant leaves may be nitrogen poor, with hard to digest plant cell wall components (Salem et al., 2017), but we didn’t find polysaccharide degrading pathways or extensive biosynthesis of essential amino acids in the Lv-StB genome. Therefore, the *lga* BGC is the oldest remaining feature that potentially increases host fitness. Third, the reduced coding density seen in the Lv-StB genome may be indicative of a recent transitional event (Lo et al., 2016), such as strict host association or a move to vertical transmission. Fourth, even though we found genes missing from all DNA repair pathways, which is thought to be a driver for increased AT content in symbiont genomes (McCutcheon and Moran, 2012), and increase in AT may have an adaptive component that reduces the metabolic costs of symbionts (Dietel et al., 2019), the GC content of the Lv-StB genome is not very different from free-living *B. gladioli* strains, when compared to other “transitional” symbionts. For instance, “*Candidatus* Pantoea carbekii” has a reduced genome and like Lv-StB lacks full-length DNA polymerase I (Kenyon et al., 2015). This strain has a GC content almost 30% lower than its closest freeliving relatives (Lo et al., 2016), while the genome of Lv-StB has a GC content ∼10% lower than its closest relatives. Therefore, we propose that loss of DNA repair pathways and other genome degradation events in Lv-StB occurred very recently, after the acquisition of *lga*.

It can be envisioned that *lga* provided a sustained survival advantage in an environment where lagriamide consistently reduced egg fungal infections, and there was positive selection on beetles that vertically transmitted *lga*-bearing symbionts. In *L. villosa* symbionts are stored extracellularly, and they are spread onto the outside of eggs as they are laid. According to observations in the congeneric species *L. hirta*, the symbionts first enter through the egg micropyle to reach the embryonic organs where they are housed throughout larval development (Jürgen Stammer, 1929). It is thus likely that only a few of the cells are vertically transmitted by colonizing these structures, potentially providing the population bottlenecks that could have caused initial accumulation of deleterious mutations that started the process of genome reduction. Meanwhile, loss of certain proteins limiting growth rate (see above) may have been selected through increased Lv-StB populations and compound production. It is unknown to what extent Lv-StB is genetically isolated in the larval or adult host, but we found evidence of ongoing horizontal transfer events in the recent past, presumably through contact with a complex microbiome associated with *L. villosa* egg surfaces. These horizontal gene transfers likely happened concurrently with the ongoing genome reduction process and may have been compensatory for gene losses (see above). There is some precedent for extracellular symbionts with profoundly reduced genomes (Kaiwa et al., 2014; Kikuchi et al., 2009; Nikoh et al., 2011; Salem et al., 2017). For instance, the leaf beetle *Cassida rubiginosa* harbors a symbiont with the smallest genome of any extracellular organism (0.27 Mbp), “*Candidatus* Stammera capleta”, which provides pectinolytic enzymes to help break down the host’s leafy diet (Salem et al., 2017). In many of these cases, symbionts are stored as isolated monocultures within specialized structures in adult hosts, while vertical transmission is assisted by packaging symbiont cells in protective “caplets” attached to eggs (Salem et al., 2017), a “symbiont capsule” encased in chitin (Nikoh et al., 2011), or secreted in a galactan-based jelly ingested by hatched larvae (Kaiwa et al., 2014), although reduced-genome symbionts have also been known to be vertically transmitted by simple egg surface contamination (Kikuchi et al., 2009), as in *L. villosa*. Because these examples are advanced cases of genome reduction, it would be difficult to determine whether horizontal transfer events occurred before or during genome reduction, and none have been noted. The complexity of the *L. villosa* microbiome appears to be different, as it afforded ample opportunity for horizontal gene transfer even while the genome of Lv-StB was actively undergoing reduction.

Horizontal acquisition of genes has been observed in two types of reduced genome symbiont, eukaryotic parasites in the genus *Encephalitozoon* (Pombert et al., 2012), and Acetobacteraceae strains associated with the gut community of red carpenter ants (Brown and Wernegreen, 2019). In both these cases symbiont genomes were similar in size to Lv-StB (∼2 Mbp), but with far greater coding density and fewer pseudogenes. Furthermore, both *Encephalitozoon* and Acetobacteraceae strains were culturable, suggesting that they are facultative symbionts in a less advanced state of genome reduction compared to Lv-StB. The genome of Lv-StB appears to be different from these examples, because there is evidence of recent horizontal transfers, even as genes required for homologous recombination are currently missing. Either the loss of homologous recombination was very recent, or such transfers could have occurred in a RecA-independent manner. For example, plasmids could have been transferred into Lv-StB cells, followed by the RecA-independent transposition of genes to the chromosome (Harmer and Hall, 2016; Zupancic et al., 1983).

It is unclear whether Lv-StB will continue on the path of genome reduction to become drastically reduced with a <1 Mbp genome. Where symbionts are required for host survival and are genetically isolated within host cells or specialized structures, such a process appears to be irreversible and unstoppable (Bennett and Moran, 2015; Moran, 1996). However, a number of alternate fates could be envisioned for Lv-StB. With a complex microbiome, if ongoing gene losses in Lv-StB reduce its fitness past a certain point, then it could be replaced by another strain, potentially accompanied by horizontal transfer of the *lga* pathway to a less reduced genomic chassis. Alternatively, horizontal transfers of genes to Lv-StB could lead to an equilibrium of gene loss and gain. Interestingly, it appears that up until the present time horizontal transfer has not occurred fast enough to prevent widespread loss of metabolism and DNA repair in the Lv-StB genome. The host could also evolve strategies to maintain an increasingly genome-reduced Lv-StB, perhaps by selective extracellular partitioning and packaging for vertical transfer similar to the examples outlined above. However, it is unclear whether such an evolutionary path would be favorable, given that the co-infection of multiple BGC-bearing symbiont strains could be advantageous in environments with variable pathogen pressures.

In summary, evidence gathered here suggests that the introduction of the lagriamide BGC initiated genome erosion of Lv-StB, potentially through selection of beetles that transferred the symbiont vertically, leading to a population structure with frequent bottlenecks. Simultaneous advantageous gene acquisitions may have enabled the preferential survival of Lv-StB and its dominance in the adult host and the egg surface.

## MATERIALS AND METHODS

### Sequencing and assembly of *Burkholderia gladioli* Lv-StA genome

Genome sequencing of the isolated *B. gladioli* Lv-StA strain was carried out using PacBio with Single Molecule, Real-Time (SMRT) technology. For *de novo* assembly (carried out by Eurofins Genomics), the HGAP pipeline was used (Heirarchical Genome Assembly Process). Briefly, a preassembly of long and accurate sequences was generated by mapping filtered subreads to so-called seed reads. Subsequently, the Celera assembler was used to generate a draft assembly using multi-kb long reads, which in this case rendered full genome closure. Finally, the Quiver algorithm was used to correct inDel and substitution errors by considering the quality values from the bas.h5 files.

### Metagenomic binning and annotation

Metagenomic assembly files were clustered into putative genomic bins using Autometa (Master branch - commit bbcea30) (Miller et al., 2019). Contigs with lengths smaller than 3,000bp were excluded from the binning process and a taxonomy table was produced. Contigs classified as bacterial were further binned into putative genomic bins using run_autometa.py. Unclustered contigs were recruited into clusters using ML_recruitment.py. Results were summarized using cluster_process.py. Resultant genome bins were compared to earlier versions (Flórez et al., 2018) using Mash version 2.1.1 (Ondov et al., 2016) which hashes genomes to patterns of *k*-mers (sketching) allowing for rapid distance calculations between two sketches. All bins were sketched and distances were computed in a pairwise fashion. Pairwise distances were visualized in R as a dendrogram and enabled the determination of equivalent old and updated putative genome bins between analyses. The updated putative genomic bins were annotated using Prokka version 1.13 (Seemann, 2014), with genbank compliance enabled. Reference genomes downloaded from NCBI were similarly annotated with Prokka in order to maintain consistency between datasets. Amino acid sequences of open-reading frames (ORFs) were further annotated using DIAMOND blastp version 0.9.21.122 (Buchfink et al., 2015) against the diamond formatted NR database. The search was limited to returning a maximum of 1 target sequence and the maximum number of high-scoring pairs per subject sequence was set to 1. Results were summarized in BLAST tabular format with qseqid (Query gene ID), stitle (aligned gene ID), pident (Percentage of identical matches), evalue (Expected value), qlen (Query sequence length) and slen (aligned gene sequence length) as desired parameter output. Pseudogenes were identified by finding Lv-StB genes that were more than 20% shorter than their respective BLAST matches. This criteria has been used previously to identify pseudogenes (Kwan and Schmidt, 2013; Lerat and Ochman, 2005).

Coding density was calculated as the sum of all protein coding sequences (coding sequence) as a percentage of the sum of all contigs (total sequence). In cases where protein coding genes were found to overlap, the length of the overlap region was counted only once. This calculation was performed for all binned genomes, on both initial datasets (genbank files generated during Prokka annotations) and edited datasets where pseudogenes had been removed. For the identification and count of genes encoding transposases and hypothetical proteins, protein-coding gene amino acid files (*.faa) containing Prokka annotations were parsed for gene descriptions containing “transposase” and “hypothetical” strings.

### Taxonomic classification of genome bins

Putative genome bins clustered from the *L. villosa* metagenomic dataset were taxonomically classified using GTDB-Tk v0.2.2 (reference database gtdbtk.r86_v2) with default parameters (Parks et al., 2018). GTDB-Tk identifies and aligns 120 bacterial marker genes per genome before calculating the optimal placement of the respective alignments in the pre-computed GTDB-Tk reference tree which consists of 94,759 genomes (Dataset S1B). A species was assigned to a genome if it shared 95% or more ANI with a reference genome.

### Identification of “core” genes

The set of “core” genes generally found in even the most reduced symbiont genomes was taken from Table 2 of McCutcheon and Moran 2012 (McCutcheon and Moran, 2012). GFF files produced for *B. gladioli* Lv-StA and the metagenomic bins by Prokka were searched for the following gene symbols: ‘*dnaE*’, ‘*dnaQ*’, ‘*rpoA*’, ‘*rpoB*’, ‘*rpoC*’, ‘*rpoD*’, ‘*groL*’, ‘*groS*’, ‘*dnaK*’, ‘*mnmA*’, ‘*mnmE*’, ‘*mnmG*’, ‘*sufS*’, ‘*sufB*’, ‘*sufC*’, ‘*iscS*’, ‘*iscA*’, ‘*iscU*’, ‘*rluA*’, ‘*rluB*’, ‘*rluC*’, ‘*rluD*’, ‘*rluE*’, ‘*rluF*’, ‘*infA*’, ‘*infB*’, ‘*infC*’, ‘*fusA*’, ‘*tsf*’, ‘*prfA*’, ‘*prfB*’, ‘*frr*’, ‘*def*’, ‘*alaS*’, ‘*gltX*’, ‘*glyQ*’, ‘*ileS*’, ‘*metG*’, ‘*pheS*’, ‘*trpS*’, ‘*valS*’, ‘*rpsA*’, ‘*rpsB*’, ‘*rpsC*’, ‘*rpsD*’, ‘*rpsE*’, ‘*rpsG*’, ‘*rpsH*’, ‘*rpsI*’, ‘*rpsJ*’, ‘*rpsK*’, ‘*rpsL*’, ‘*rpsM*’, ‘*rpsN*’, ‘*rpsP*’, ‘*rpsQ*’, ‘*rpsR*’, ‘*rpsS*’, ‘*rplB*’, ‘*rplC*’, ‘*rplD*’, ‘*rplE*’, ‘*rplF*’, ‘*rplK*’, ‘*rplM*’, ‘*rplN*’, ‘*rplO*’, ‘*rplP*’, ‘*rplT*’, ‘*rplV*’, ‘*rpmA*’, ‘*rpmB*’, ‘*rpmG*’, ‘*rpmJ*’, ‘*tRNA-Met*’, ‘*tRNA-Gly*’, ‘*tRNA-Cys*’, ‘*tRNA-Phe*’, ‘*tRNA-Lys*’, ‘*tRNA-Ala*’, ‘*tRNA-Glu*’, ‘*tRNA-Pro*’, ‘*tRNA-Gln*’, ‘*tRNA-Ile*’. The presence of single or multiple examples of these genes per genome/bin was tabulated in excel to produce Dataset S1D, and the percentage of the core gene set found in a genome/bin was used for Table 1.

### Annotation and analysis of BGCs

Putative biosynthetic gene clusters were identified in all binned genomes using the AntiSMASH (Blin et al., 2019) docker image (Image ID: 8942d142d9ac). Entire genome HMMer analysis was enabled, and identified clusters were compared to both antiSMASH-predicted clusters, the MIBiG database and secondary metabolite orthologous groups. Similarities between identified putative biosynthetic gene clusters were assessed using BiG-SCAPE version 20181005 (Navarro-Muñoz et al., 2018) in “glocal” mode.

### Construction of multilocus species tree

Genomes of *Burkholderia* Lv-StB, *B. gladioli* Lv-StA and binned genomes taxonomically classified within the *Burkholderia* genus: DBSCAN_round6_14, DBSCAN_round6_18 and DBSCAN_round4_0, were uploaded to the AutoMLST website (Alanjary et al., 2019). A concatenated species tree was constructed in *de novo* mode, with default options as well as the IQ-TREE Ultrafast Bootstrap analysis and ModelFinder options enabled.

### Calculation of average nucleotide identities

The average nucleotide identities (ANIs) of *Burkholderia* Lv-StB and *B. gladioli* were calculated in a pairwise manner using FastANI (Jain et al., 2018) against 45 Burkholderia reference genomes downloaded from NCBI. A total of 13 genomes shared over 85% ANI (Dataset S1H). These genomes were all identified as *B. gladioli* species and were used in downstream analyses.

### Quantification of divergence between *Burkholderia* Lv-StB and *B. gladioli* A1

Orthologous protein sequences were identified in non-pseudogene sequence files of *Burkholderia* Lv-StB and 13 closely related genomes (identified through the ANI analyses: *B. gladioli* Lv-StA, *B. gladioli* A1, *B. gladioli* UCDUG, *B. gladioli* FDAARGOS_389, *B. gladioli* ATCC25417, *B. gladioli* Co14, *B. gladioli* SN82F6, *B. gladioli* ATCC10248, *B. gladioli* NBRC13700, *B. gladioli* FDAARGOS_188, *B. gladioli* MSMB1756, *B. gladioli* BSR3, *B. gladioli* KACC11889) using OMA version 2.2.0 (Altenhoff et al., 2018). A subset (797 groups) of the resultant orthologous groups (OGs) was identified which included genes from all 14 genomes used in the analysis. Each set of OG sequences were aligned using MUSCLE v3.8.31 (Edgar, 2004) and corresponding nucleotide files were extracted and aligned against the amino acid sequences using the PAL2NAL docker image (Image ID: ce3b1d7d83ab) (Suyama et al., 2006) using codon table 11 and specifying no gaps with paml as the output format. The resultant paml files were used to estimate pairwise dS (synonymous divergence rate), dN (non-synonymous divergence rate) and kappa (transition/transversion ratio) between individual genes per orthologous group with codeml (Yang, 2007) in the PAL2NAL package. The following parameters were specified in the control file: runmode = -2 (pairwise), model = 0 (one) fix_kappa = 0 (kappa to be estimated), fix_omega = 0 (estimate omega) where omega is the dN/dS ratio, with initial omega set to 0.2. Any orthologous gene sets that included genes that gave a dS value over 3 were removed from the analysis (Yang, 2014). Individual sequences from remaining OGs were then gathered into genome-specific files (i.e all Lv-StB genes in all OGs were moved into an ordered Lv-StB.faa/.ffn file). Stop codons were removed from each nucleotide sequence. Sequences per genome were then concatenated to produce a single sequence per genome. The concatenated amino acid sequences and corresponding nucleotide sequences were aligned against one another using PAL2NAL as performed for individual genes. Pairwise estimations of dS, dN and kappa were calculated as before using codeml. Additionally, the concatenated genes were analysed a second time using an alternative control file, in which the model was set to 2. The likelihood ratio test value between pairwise null and alternative likelihood scores was calculated (2x Alt_lnl - Null_lnl) for Lv-StB relative to the 13 reference genomes and found to be 0 in all cases indicating that the omega (dN/dS) ratio was consistent between Lv-StB and the reference genomes. Individual dS values were used to estimate divergence between Lv-StB and the 13 reference genomes using divergence rates estimated by Silva and Santos-Garcia (Silva and Santos-Garcia, 2015) (Table S1) in the equation: Age of divergence (Mya) = dS ÷ divergence rate x 1,000,000. As Lv-StB shared the greatest ANI with *B. gladioli* A1, the kappa value found between these two genomes (7.03487) was used for amelioration estimates.

### Pangenome analysis

To assess the pangenome of Lv-StB and other *B. gladioli* genomes, GFF files generated by Prokka were analysed using Roary (Page et al., 2015) which identifies core and accessory genes per genome. Concatenation and alignment of orthologous genes was enabled in Roary and used to build a phylogenetic tree with FastTree version 2.1.10 (Price et al., 2010). The resultant phylogenetic tree and presence/absence matrix of genes in all genomes were visualized with the roary_plots.py script. Additionally, non-pseudogenes of all genomes were annotated against the KEGG database (Kanehisa et al., 2019; Kanehisa and Goto, 2000) using kofamscan (Aramaki et al., 2019) with output in mapper format. An overview of the completeness of general metabolic pathways was visualized using KEGG-Decoder (Graham et al., 2018) with kofamscan annotations. For specific pathways of interest (amino acids, DNA repair, nucleotide de novo biosynthesis), presence/absence matrices of genes per KEGG pathway entry were visualized in R version 3.6.0 using the tidyr, ggplot2 and viridis libraries.

### Identification of genes putatively acquired by horizontal transfer

All amino acid sequence files of non-pseudogenes of all genomes used in the ANI analysis (*B. gladioli* Lv-StA and 45 *Burkholderia* reference genomes) were concatenated and converted into a DIAMOND BLAST (Buchfink et al., 2015) database (build 125). The amino acid sequence files of non-pseudogenes in Lv-StB were then searched against this database. All genes that had no significant hit, or a significant hit but with a shared percentage of less than 50% were considered “unique” to Lv-StB. Non-pseudogenes of Lv-StB that were not found to have an ortholog in the OMA analysis were used to validate this list. These “unique” genes were then compared to the NR database using DIAMOND blastp (as described above) and any genes that had no significant hit were removed from the “unique” set of genes. Manual inspection of the remaining genes resulted in the removal of any genes that were closely related to *B. gladioli* genes (i.e. found within *B. gladioli* genomes other than the ones investigated here). The remaining unique genes were henceforth considered as gene potentially acquired via horizontal transfer. This list was expanded with other genes within Lv-StB, for which homologs in *Burkholderia* genomes could be found, but the closest match against the NR database belonged to a different genus. For example, E5299_02249 of the “addic” group shared 53.1% sequence identity with a gene from *B. contaminans* strain LMG 23361 but shares a higher sequence identity of 96.9% with *Ochrobactrum pituitosum*.

### De-amelioration of putatively horizontally transferred genes

The method of Lawrence and Ochman (Lawrence and Ochman, 1997) was implemented in Python and is available at https://bitbucket.org/jason_c_kwan/age_horizontal_transfers.py. The script takes as input 1. an in-frame nucleotide FASTA file containing the sequences of putatively horizontally transferred genes, 2. an in-frame nucleotide FASTA file containing a comparison set of gene sequences from the genome, 3. a synonymous mutation rate in substitutions per 100 sites per million years, 4. a nonsynonymous mutation rate in substitutions per 100 sites per million years, 5. a transition/transversion ratio (κ), 6. a step time in millions of years, and 7. a maximum time to iterate to. The script outputs GC content of each codon position (plus overall GC) at each timepoint, and reports the estimated age of the gene cluster as corresponding to the iteration with the smallest sum of squared deviations from equations 2–4. The substitution rates used in our calculations were half of the divergence rates estimated by Silva and Santos-Garcia (Silva and Santos-Garcia, 2015), BAU: synonymous 0.55, nonsynonymous 0.05; BOB: synonymous 3.95; nonsynonymous 0.26; BPN: synonymous 3.2, nonsynonymous 0.28; BFL: synonymous 4.45, nonsynonymous 0.395. A value of 7.0348 was used for κ, previously calculated for *Burkholderia* Lv-StB and *B. gladioli* A1 (see above). A step time of 0.005 My and a maximum time of 50 My was used in all calculations. The comparison gene set included only non-pseudogenes that were not identified as putative horizontally transferred genes.

### Microbial community analysis

16S rRNA amplicon sequence datasets analysed via oligotyping previously (Flórez et al., 2018) were re-analysed using Mothur v.1.40.3 (Schloss et al., 2009). Reads shorter than 200bp, or containing ambiguous bases or homopolymeric runs longer than 7 bases were removed from the dataset. Chimeric sequences were identified using VSEARCH (Rognes et al., 2016) and removed from the dataset. Reads were taxonomically classified against the Silva database (version 132) and all reads classified as unknown, eukaryotic, mitochondrial or as chloroplasts were removed from the dataset. Reads were aligned using the Silva database (v. 132) as reference and clustered into operational taxonomic units (OTUs) at a distance of 0.03: an approximation to bacterial species. Counts of OTUs per sample were generated and the top 10 most abundant OTUs were plotted (Figure S3). The top 50 most abundant OTUs were queried against the “nt” nucleotide database using blast for taxonomic classification.

### Data availability

The complete *Burkholderia gladioli* Lv-StA genome, the draft assembly of the *Burkholderia* Lv-StB genome and other bacterial metagenomic bins will be deposited in GenBank, and the respective accession numbers will be included in the accepted version of this manuscript.

## ACKNOWLEDGMENTS

The authors wish to thank Jason Peters and Marc Chevrette (both UW-Madison) for helpful comments which improved the manuscript. The authors acknowledge funding from the Gordon and Betty Moore Foundation (MMI-6920, KLM). This research was performed in part using the computer resources and assistance of the UW-Madison Center for High Throughput Computing (CHTC) in the Department of Computer Sciences. The CHTC is supported by UW-Madison, the Advanced Computing Initiative, the Wisconsin Alumni Research Foundation, and Wisconsin Institutes for Discovery, and the National Science Foundation and is an active member of the Open Science Grid, which is supported by the National Science Foundation and the U.S. Department of Energy’s Office of Science.

---

**Table S1.**
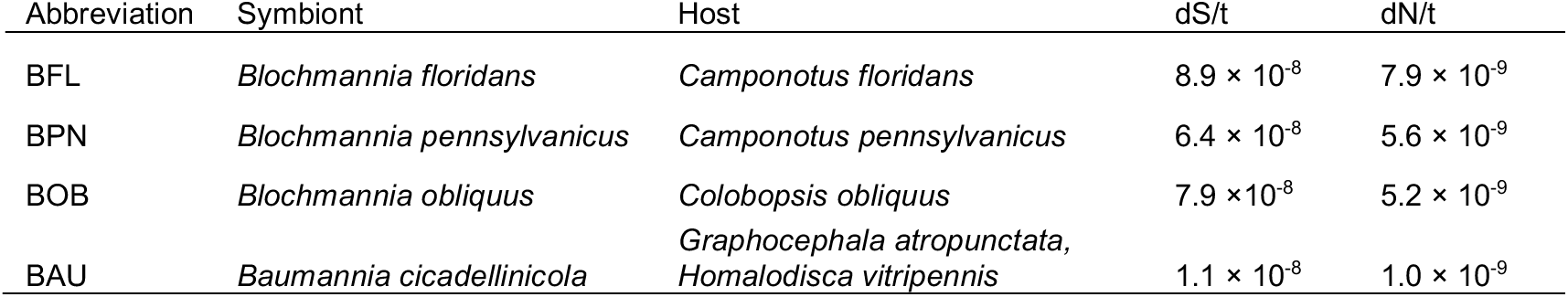
Divergence rates used in this study (taken from Silva and Santos-Garcia 2015 (Silva and Santos-Garcia, 2015)).

**Dataset S1.** Comparative analysis of the *Burkholderia* Lv-StB to other genomes in the genus *Burkholderia*.

**Figure S1.**
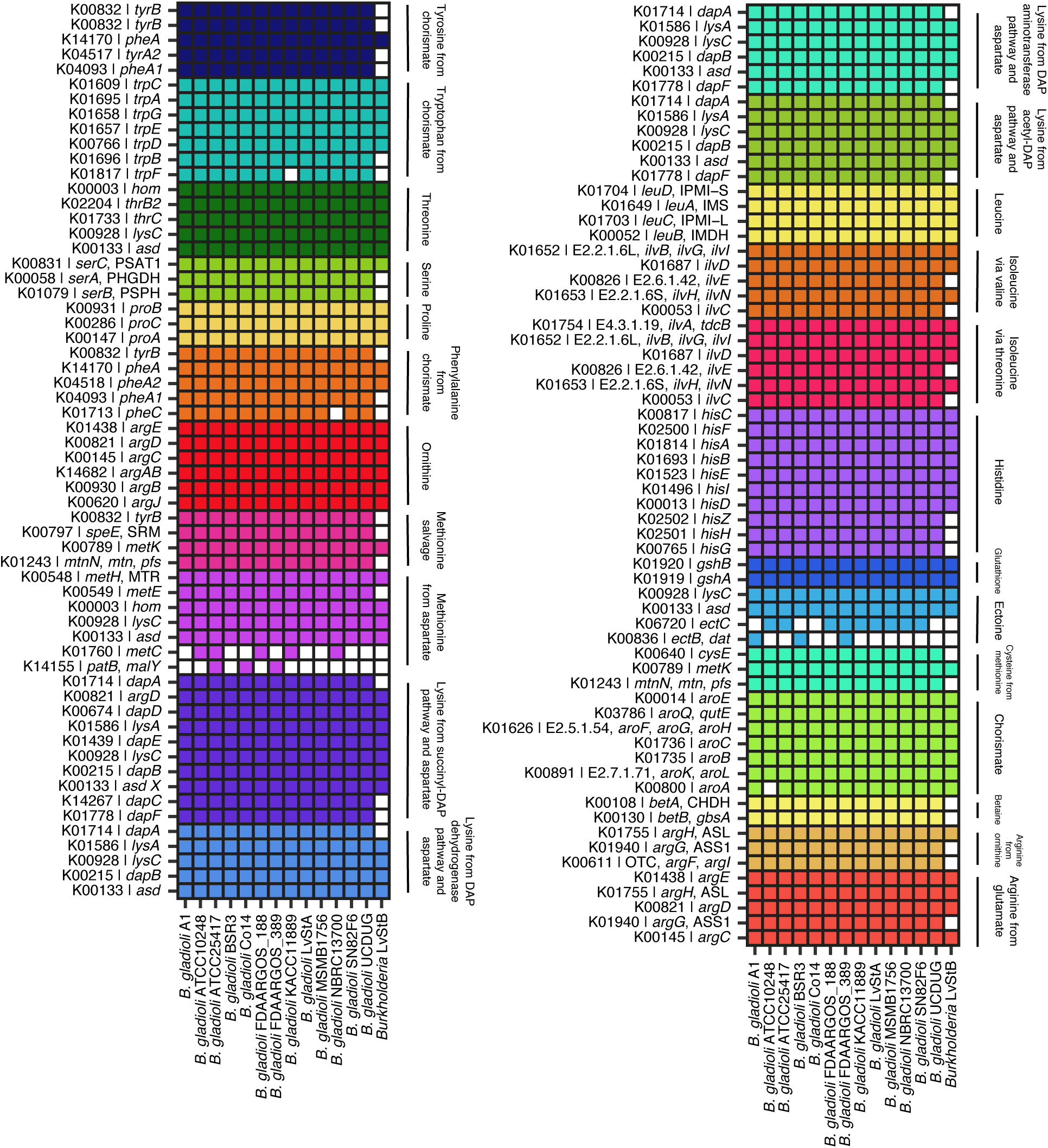
Completeness of amino acid biosynthesis pathways in *Burkholderia* Lv-StB in comparison to closely-related strains of *B. gladioli*.

**Figure S2.**
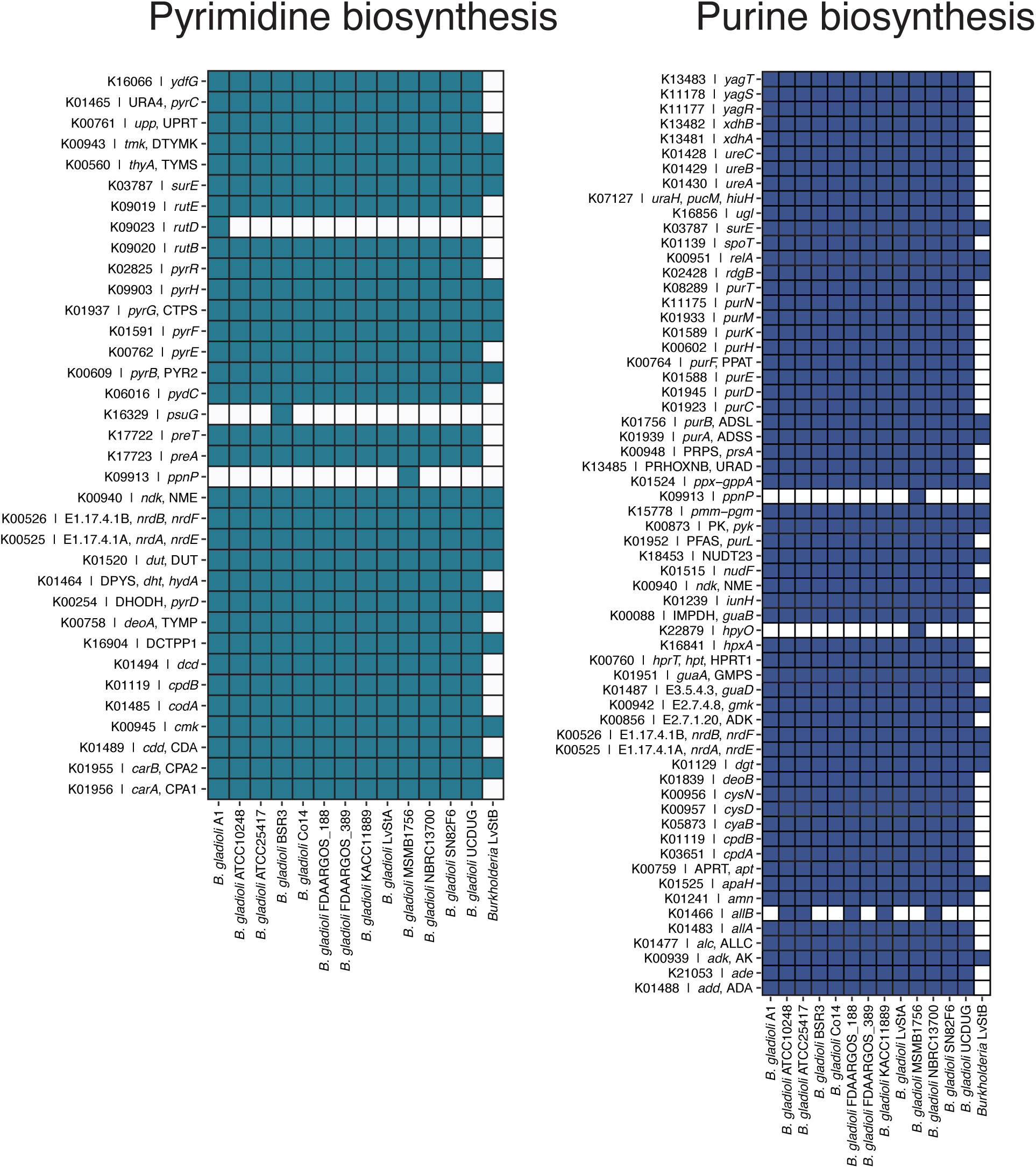
Completeness of nucleotide biosynthesis pathways in *Burkholderia* Lv-StB in comparison to closely-related strains of *B. gladioli*.

**Figure S3.**
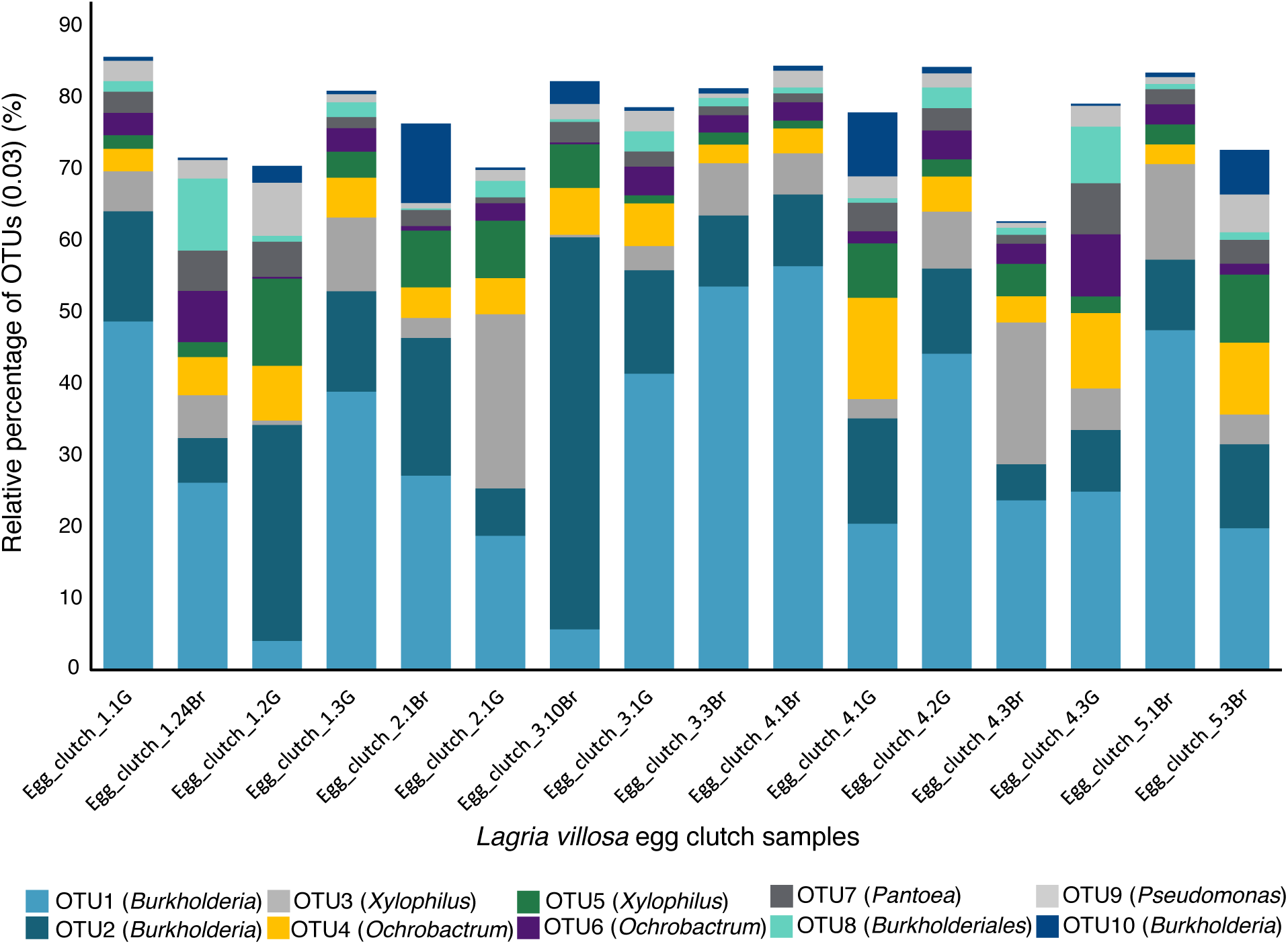
Reanalysis of 16S rRNA amplicon data used in Flórez *et al*. 2017 (Flórez et al., 2017), showing distribution of dominant microbial communities associated with *L. villosa* egg clusters. Amplicon 16S rRNA gene sequences were clustered into operational taxonomic units (OTUs) at a distance of 0.03 as an approximation to bacterial species. The putative taxonomic classification of each OTU is indicated with a colored key. Abundance of the top 10 most abundant OTUs is indicated as a relative percentage of the total reads for each cluster of eggs collected.

